# Liquid Crystalline Layering and Divalent Cations Cooperatively Enhance DNA Condensation

**DOI:** 10.1101/2023.08.29.555320

**Authors:** Sineth G. Kodikara, Prabesh Gyawali, James T. Gleeson, Antal Jakli, Samuel Sprunt, Hamza Balci

## Abstract

The layered liquid crystalline (LC) phases formed by DNA molecules which include rigid and flexible segments (‘gapped DNA’) enable the study of both end-to-end stacking and side-to-side lateral interactions that drive the condensation of DNA molecules. The resulting layer structure exhibits long-range inter-layer and intra-layer positional correlations. Using synchrotron small-angle x-ray scattering (SAXS) measurements, we investigate the impact of divalent Mg^2+^ cations on the stability of the inter- and intra-layer DNA ordering as a function of temperature between 5-65 °C and for different terminal base pairings at the blunt ends of the gapped DNA constructs, which mediate the strength of the attractive end-to-end interaction. We demonstrate that the stabilities at a fixed DNA concentration of both inter-layer and intra-layer order are significantly enhanced even at a few mM Mg^2+^ concentration. The stability continues to increase up to ∼30 mM Mg^2+^ concentration, but at higher (∼100 mM) Mg^2+^ content repulsion between positive ions counteracts and reverses the increase. On the other hand, sufficiently strong base-stacking interactions promote intra-layer order even in the absence of multivalent cations, which demonstrates the impact of liquid crystal layering on the DNA condensation process. We discuss the implications of these results in terms cation-mediated DNA-DNA attraction.

## Introduction

Condensation of DNA is an indispensable feature of biology and is prevalent in settings ranging in complexity from the highly organized eukaryotic nuclei to the tightly packed viral capsids (1). This compaction process has implications beyond just information storage and plays critical roles in several regulatory processes, such as gene expression regulation. Electrostatic interactions are at the core of this compaction process, which is facilitated by positively charged surfaces on the histone proteins and the presence of mono- and multi-valent ions in the environment (2–4).

*In vitro* studies have shown that isolated DNA molecules may condense in the presence of multivalent cations. While earlier studies demonstrated the condensation process requires ions with +3 or higher valency (such as spermidine Sd^3+^ or spermine Sm^4+^) (5), recent studies showed spontaneous condensation with +2 valency ions (such as Mg^2+^) (6), where Mg^2+^ ions bridged minor grooves of double stranded DNA (7) (which will be referred to as DNA for brevity). The spermidine and spermine polyamines are chain-molecules having extended size and, in these respects, differ significantly from the much smaller monovalent K^+^ or divalent Mg^2+^ ions. Nevertheless, these ions are all present in cellular environment(8) at comparable concentrations to those in which DNA condensation has been observed, which has motivated numerous theoretical and computational efforts to understand the condensation process in different ionic environments (9–12). The divalent ion Mg^2+^, which is the relevant ion for this investigation, has a cellular concentration of 5-30 mM, most of which is bound (e.g., to ATP). Only about ∼1 mM of the Mg^2+^ is typically free (13).

The DNA condensation process presents a challenge to mean-field Poisson-Boltzmann theory, which predicts that rod-like DNA molecules with a diffuse ion atmosphere around them do not attract each other, regardless of the valency and concentration of the cations (14, 15). The diffuse atmosphere picture is adequate to describe the association of monovalent cations with DNA (16). Divalent Mg^2+^, however, can interact with DNA in two modes: the diffuse mode where the ions (in fully-hydrated form) associate non-specifically with DNA and a localized mode where dehydrated or partially hydrated ions accumulate in specific regions of DNA (17, 18) having irregular shape, such as minor or major grooves. In concentrated DNA solutions, the attraction between DNA molecules is typically attributed to spatial correlations between the multivalent cations that localize (or are adsorbed) on one DNA molecule and the negative charges on the phosphate backbone of another molecule (19). In this DNA-ion-DNA bridging model, cations between DNA helices mediate attractive interactions. Interestingly, DNA duplexes in which adenine and thymine nucleotides were placed consecutively (AAA…TTT, called ‘homogenous’) or alternatingly (ATATAT…, called ‘heterogeneous’) showed significant differences in their condensation behavior: The homogenous construct condensed in the presence of divalent cations, while the heterogeneous construct (and the constructs with random sequence) did not (6). All-atom molecular dynamics simulations showed specific cation binding to homogeneous constructs at major grooves, whose surfaces are exposed to neighboring DNA molecules. Since a similar effect was not observed for heterogenous constructs (6), this specific binding was proposed as a mechanism for a sequence-dependent condensation process.

At sufficiently high concentration, DNA duplexes in aqueous solution also spontaneously condense into well-known chiral nematic and columnar liquid crystal (LC) mesophases (20, 21). Remarkably, LC formation may occur via an initial end-to-end stacking of ultrashort duplexes (20–25), producing longer aggregates whose aspect ratio (*L/d*, where *L* is the length and *d* the diameter) satisfies the Onsager criterion (*L/d* > 4.7) for orientational ordering of semi-rigid spherocylindrical particles (26, 27). Incorporating a flexible ‘gap’ of single-stranded DNA between two fully-paired duplexes (with *L* below the ∼50 nm persistence length) further enriches the LC phase diagram through the appearance of previously unobserved layered (smectic) phases (28–30). Two different smectic layer structures are observed: a “bilayer” stacking with ∼2 duplex periodicity, prevailing for longer gap lengths (e.g., 20 bases) and a “monolayer” stacking (∼1 duplex periodicity, occurring for gaps of 4 – 10 bases in length (29). The bilayer phase is mediated by enthalpic end-to-end attraction between duplexes and by a segregation of the flexible ‘gap’ segments into layers that becomes entropically favorable when the side-to-side separation between rigid duplex segments is reduced. In addition to interlayer ordering along the duplex axis, the bilayer smectic phase also develops hexagonal positional correlations within the layers (smectic-B phase), driven by side-to-side (‘helix-helix’) interactions at higher DNA concentration. This lateral ordering of duplexes – with characteristic signature in X-ray diffraction – provides a consistent testbed for experimentally investigating key factors that drive DNA condensation under more general (e.g., non-liquid crystalline) conditions.

Here we describe results on the effect of divalent cation concentration on the stability of smectic-B order (a layered LC phase with hexagonal positional order within the layers) in dense solutions of three “gapped” DNA (GDNA) constructs consisting of two 48-bp long rigid segments connected by a flexible, 20 nucleotide (nt) long single-stranded thymine spacer – an architecture abbreviated as 48-20T-48. The constructs have random sequences and are identical except for the single base pairs terminating the opposing blunt ends of the duplexes. On this basis, we refer to them separately as AT-AT, AT-GC, or GC-GC (Fig.1A). As we demonstrated earlier (30), 48-20T-48 GDNA constructs, suspended at ∼280 mg/ml concentration in 150 mM Na^+^ solution, exhibit bilayer smectic ordering whose stability against temperature differs significantly within the range 35-64 °C. We thereby established a clear correlation between the identity of the terminal base pairs and the thermal stability of the bilayer structure. In the present work, using small angle X-ray scattering (SAXS), we now demonstrate the effect of progressively increasing the concentration of Mg^2+^ (in the range 0-100 mM, at constant 150 mM monovalent Na^+^ concentration, Fig. 1B) on the thermal stability of the bilayer smectic-B phase of the GDNA constructs in Fig. 1A. The bilayer structure in Fig. 1B is rendered with “folded” duplexes based on evidence for this morphology from experiment (29) and MD simulations (31) for constructs with sufficiently long “gap”, e.g., 20 T in the present case. In addition to demonstrating the impact of end-to-end stacking and entropic effects on generating an effective condensation process at modest DNA concentrations (compared to the range explored in earlier studies), we report significant variation in the stability of LC phases even at low mM, physiologically relevant Mg^2+^ concentration, illustrating the significance of divalent cation concentrations on the DNA condensation process and confirming that the presence of higher valency ions is not a fundamental requirement for sequence-independent DNA condensation.

**Figure 1.**
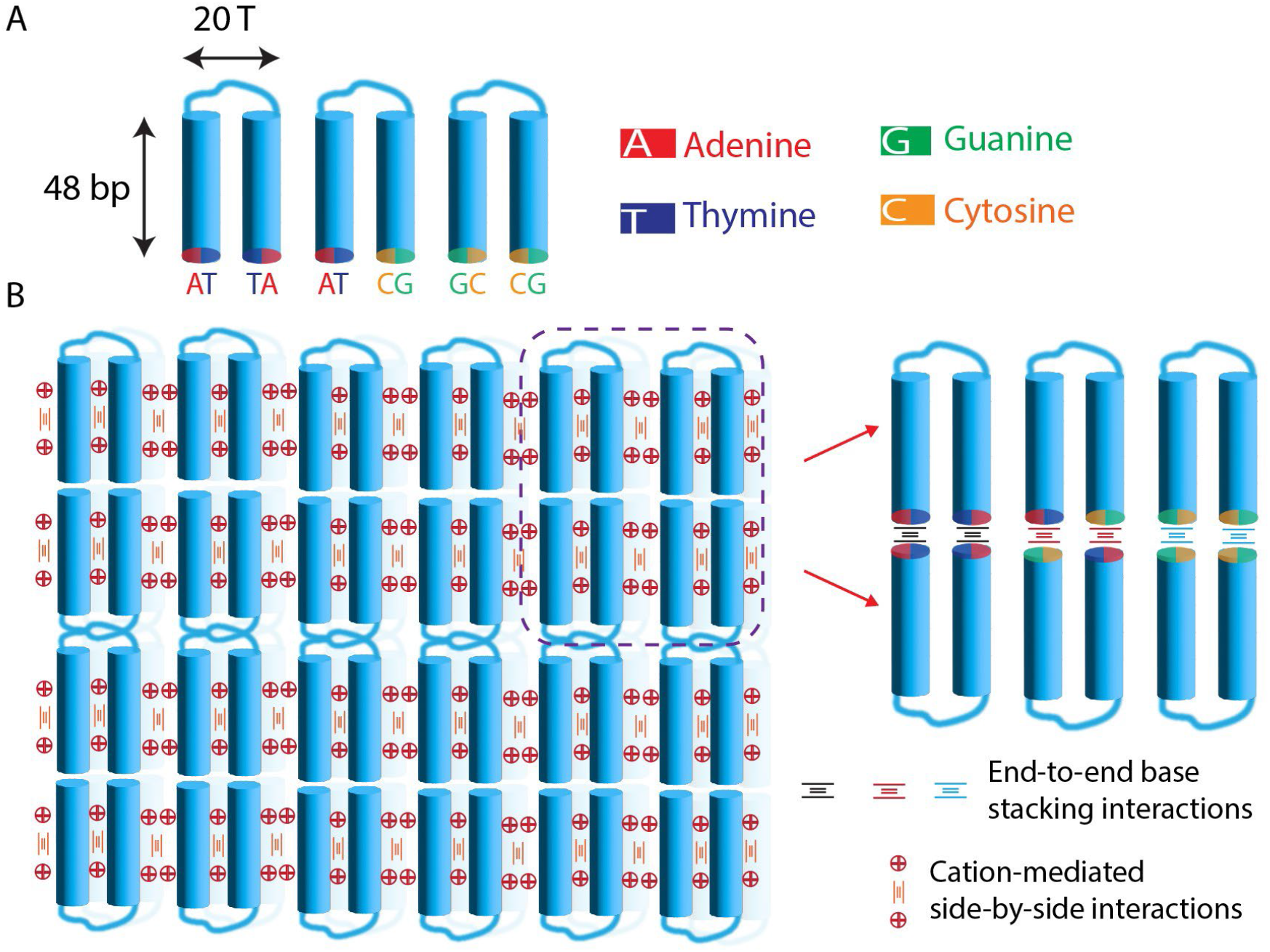
Schematic motifs of 48-20T-48 GDNA constructs. A) The AT-AT, AT-GC, and GC-GC constructs are identical except for their terminal base pairs (used to name the constructs). B) The bilayer phase of GDNA constructs where base stacking and side-by-side (helix-helix) interactions are indicated.

## Results

We conducted SAXS experiments at beamline 11-BM at the National Synchrotron Light Source II. A typical SAXS pattern from smectic GDNA samples features sharp, small-angle peaks representing diffraction from the stacking of duplexes along the layer normal, together with a diffuse or sharp peak at a wider angle that corresponds to short-or long-range positional correlations between duplexes within the layers – i.e., smectic-A or smectic-B phase, respectively. Fig. 2 displays azimuthally averaged SAXS intensity vs scattering wavenumber *q* as a function of temperature for 260-265 mg/ml solutions of the AT-AT, AT-GC, and GC-GC constructs, each at 0 mM, 2 mM, 30 mM or 100 mM Mg^2+^ ion concentration and fixed 150 mM Na^+^ concentration. For each sample, the fundamental order of diffraction observed at small angle is centered on wavenumber *q*_*1*_ = 0.182 nm^-1^ (with ±0.002 nm^-1^ variation across different constructs) at the lowest temperature (5-7 °C), which gives a smectic layer spacing of *d = 2π/q*_*1*_ = 34.5 nm (±0.4 nm). A weaker first harmonic appears at *2q*_*l*_. The layer spacing slightly exceeds the 32 nm length of two 48-bp duplexes and confirms a bilayer structure(29, 30, 32). As previously established from SAXS studies (32), the sharp wider angle peak at *q*_*w*_ = 2.15 nm^-1^ (with ±0.02 nm^-1^ variation across different constructs), observed in a significant subset of samples in Fig. 2, represents the lowest order diffraction from hexagonal positional ordering of the duplexes within the layers that characterizes the smectic-B phase. The intralayer center-to-center spacing between duplexes is *a = 4π/*√3*q*_*w*_ = 3.37 nm (±0.03 nm) at 5-7 °C.

**Figure 2.**
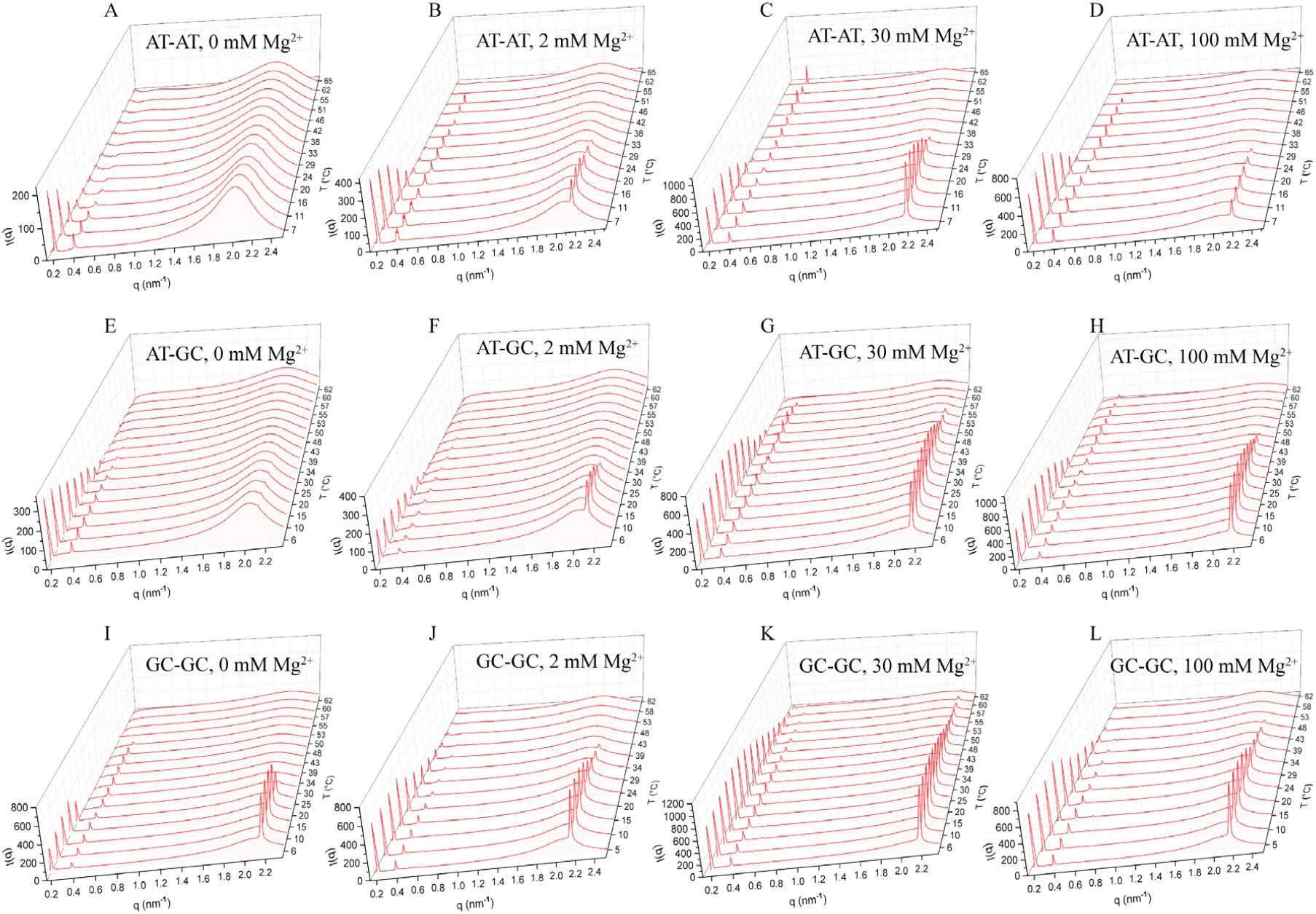
Temperature dependence (on heating from ∼5-65 °C) of the azimuthally-averaged SAXS intensity vs. scattering wave number *q* for the GDNA samples studied. A-D) AT-AT samples, E-H) AT-GC samples, I-L) GC-GC samples at 0, 2, 30, and 100 mM Mg^2+^ concentration, respectively. All samples were maintained at 260-265 mg/ml DNA and 150 mM Na^+^ concentration.

Figs. 2A-D present temperature-dependent SAXS data for AT-AT samples at 0, 2, 30 and 100 mM Mg^2+^, respectively. The diffuse wider-angle peak for the 0 mM Mg^2+^ sample (Fig. 2A) signifies a smectic-A phase with short-range positional order in the layers. When 2 mM Mg^2+^ is introduced (Fig. 2B), a sharp wider-angle peak appears at *q*_*w*_ and lower temperatures, indicating a cation-driven condensation of the duplexes into an ordered structure within the layers (smectic-B phase). The peak height decreases with increasing temperature, and the peak disappears around 29 °C. The addition of divalent salt also stabilizes the bilayer stacking (identified by the small angle peak at *q*_*1*_ = 0.182 nm^1^) to higher temperature than in the 0 mM Mg^2+^ sample. Increasing the Mg^2+^ concentration to 30 mM (Fig. 2C) enhances the thermal stability of both the inter- and intra-layer order, expanding the smectic-B range to higher temperature. However, with further increase in Mg^2+^ concentration (to 100 mM, Fig. 2D), the wider-angle peak intensity, as well as the bilayer melting temperature, decreases, indicating that the effective helix-helix attraction achieved at 30 mM begins to weaken at higher Mg^2+^ concentration. Interestingly, in the samples containing Mg^2+^ (Fig. 2B-D), a weak but sharp, small-angle peak at 0.359 nm^-1^ remains after the bilayer smectic-B melts and contributes up to the highest temperature (65 °C) studied. This peak corresponds to a layer spacing of ∼17.5 nm, slightly larger than the length of a single duplex. Careful examination of the data in Fig. 2 reveals that this peak develops from a splitting of the first harmonic of the bilayer peak (*q* ∼ 0.359 nm^-1^) at elevated temperature, consistent with the bilayer smectic-B to “monolayer” smectic-A structural transition that we previously identified (32) in AT-terminated 48-10T-48 samples without divalent cations but at significantly higher (∼280 mg/ml) DNA concentration.

Figs. 2E-H show SAXS data for AT-GC samples at 0, 2, 30 and 100 mM Mg^2+^ concentration, respectively. The diffraction from the 0 mM Mg^2+^ sample (Fig. 2E) is again consistent with a bilayer smectic-A phase; however, the bilayer peak survives to a higher temperature than in the corresponding AT-AT sample, confirming the end-to-end stacking interactions are stronger in AT-GC sample compared to AT-AT sample, in agreement with published reports(30, 33–36). Also similar to AT-AT samples, the introduction of divalent cations induces lateral positional ordering of the duplexes and enhances its thermal stability up to 30 mM Mg^2+^ concentration and 53°C temperature, with a reversal in this trend developing between 30 and 100 mM (Fig. 2F-H).

Figs. 2I-L display SAXS data for GC-GC samples at 0, 2, 30 and 100 mM Mg^2+^ concentration, respectively. In this case, the 0 mM Mg^2+^ sample (Fig. 2I) exhibits a sharp *q*_*w*_ peak, indicating the existence of well-defined lateral ordering of the duplexes even in the absence of divalent cations. The same pattern of enhanced stability of the intra-layer order against temperature at 2 and 30 mM, followed by a weakening at 100 mM, is observed in this sample (Fig. 2J-L) as for the AT-AT or AT-GC constructs. However, at each cation concentration, the smectic-B ordering of the GC-GC construct is the most stable against temperature, surviving up to ∼60 °C in the 30 mM Mg^2+^ sample.

Fig. 3 shows results of an analysis used to quantify the thermal stability of bilayer (left column, Fig. 3A, C, E) and intralayer (right column, Fig. 3B, D, F) positional order based on the SAXS data in Fig. 2. A description of this analysis has been published elsewhere(30), and is summarized in the Materials and Methods section. The dashed and solid lines are fits of the data to single and double Hill functions(37, 38), respectively. The double-Hill function (resulting in characteristic melting temperatures *T*_*m1*_ and *T*_*m2*_) provides a better description of the data than the single-Hill function (resulting in a single *T*_*m*_), although in several cases data are clearly lacking in the temperature range where the “kink” in the double-Hill fit occurs. The analysis with two Hill functions implies a multi-step melting process, whose possible mechanism we discussed previously(30) for the bilayer structure sketched in Fig. 1B.

**Figure 3.**
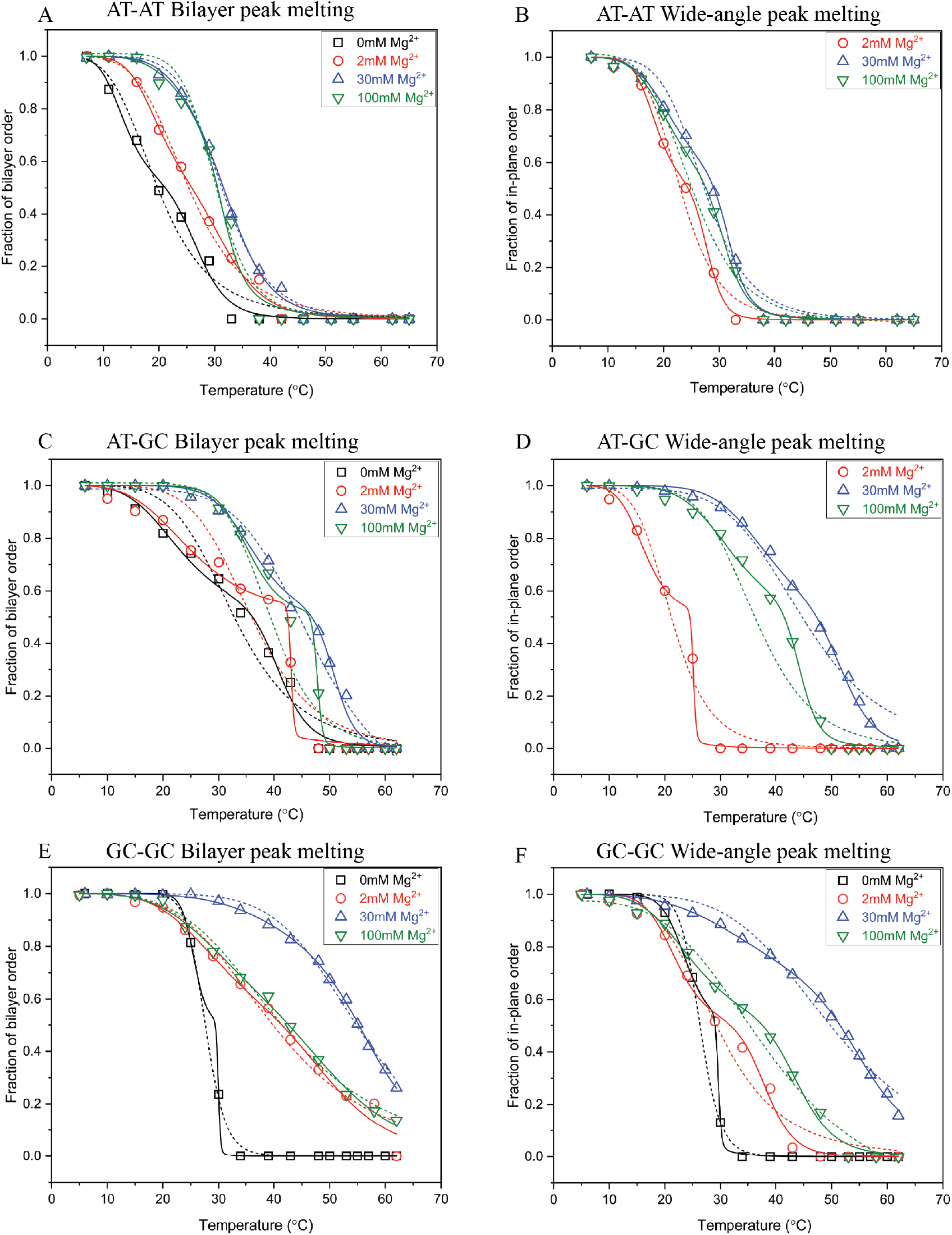
Thermal melting analysis of the SAXS data (presented in Fig. 2). A, C, E) Temperature dependence of the normalized square root of SAXS intensity integrated over first and third order small angle peaks (associated purely with scattering from stacked bilayers) for AT-AT, AT-GC, and GC-GC samples at different Mg^2+^ concentrations. B, D, F) Temperature dependence of the normalized square root of intensity integrated over the sharp wider-angle peak (associated with intra-layer positional order) for same samples. The dashed and solid lines represent single and double-Hill function fit, respectively. In a few cases (such as 2 mM Mg^2+^ in (C) and (D), or 0 mM Mg^2+^ in (E)), the fits appear to have sharp kinks at positions where we do not have data points. This potential artefact in the fits could not be avoided due to limited temperature resolution of the data.

Table 1 lists the determined values of the parameters *T*_*m*_, *T*_*m1*_, and *T*_*m2*_. (The AT-AT and AT-GC samples at 0 mM Mg^2+^ do not show sharp, wider-angle peaks, so these are not included in the table.) The intra-layer melting temperatures are consistently lower than those for the bilayer melting for each construct. This indicates that the in-layer order melts during heating before end-to-end stacking interactions are broken, again in agreement with our previous results on solutions containing monovalent cations only(30). For each construct studied, both the bilayer and intra-layer melting temperatures increase with the addition of 2 and 30 mM Mg^2+^ but then decrease at 100 mM Mg^2+^ concentration. This pattern is summarized in Fig. 4.

**Table 1:** Results for melting temperature *T*_*m*_ and for *T*_*m*1_, *T*_*m*2_ from single and double Hill function fits, respectively.

| GDNA | $\text{Mg}^{2+}$<br>(mM) | Bilayer Order | | | In-plane Order | | |
| --- | --- | --- | --- | --- | --- | --- | --- |
|  |  | Single Hill | Double Hill |  | Single Hill | Double Hill |  |
| | | $T_m$ ($^{\circ}\text{C}$) | $T_{m1}$ ($^{\circ}\text{C}$) | $T_{m2}$ ($^{\circ}\text{C}$) | $T_m$ ($^{\circ}\text{C}$) | $T_{m1}$ ($^{\circ}\text{C}$) | $T_{m2}$ ($^{\circ}\text{C}$) |
| AT-AT | 0 | 19.8±0.6 | 13.8±0.6 | 26.9±0.6 | N.A. | N.A. | N.A. |
| | 2 | 25.3±0.4 | 19.7±0.5 | 32.1±0.6 | 22.9±0.4 | 18.5±0.2 | 27.9±0.3 |
| | 30 | 31.3±0.2 | 28.2±0.6 | 33.9±0.5 | 27.3±0.5 | 21.9±0.4 | 31.8±0.4 |
| | 100 | 30.1±0.4 | 27.6±0.9 | 31.6±0.3 | 25.1±0.3 | 20.8±0.3 | 30.9±0.4 |
| AT-GC | 0 | 32.8±0.9 | 23.3±0.8 | 41.0±0.7 | N.A. | N.A. | N.A. |
| | 2 | 35.9±0.8 | 25.2±0.7 | 43.1±0.6 | 21.2±0.4 | 16.4±0.2 | 25.1±0.3 |
| | 30 | 47.1±0.7 | 36.2±0.7 | 50.9±0.4 | 44.7±0.7 | 38.1±0.5 | 51.9±0.5 |
| | 100 | 39.3±0.4 | 35.5±0.5 | 47.8±0.1 | 36.3±0.3 | 20.8±0.3 | 30.9±0.4 |
| GC-GC | 0 | 27.6±0.1 | 25.6±0.1 | 29.9±0.7 | 26.3±0.1 | 23.7±0.1 | 29.6±0.3 |
| | 2 | 39.8±0.8 | 29.8±0.9 | 50.2±0.8 | 29.3±0.8 | 22.0±0.6 | 38.3±0.6 |
| | 30 | 54.8±0.3 | 50.3±0.9 | 57.3±0.7 | 49.8±0.6 | 40.4±0.7 | 55.5±0.5 |
| | 100 | 41.5±0.5 | 31.8±0.9 | 50.9±0.8 | 39.1±0.3 | 24.8±0.6 | 44.1±0.6 |

**Figure 4.**
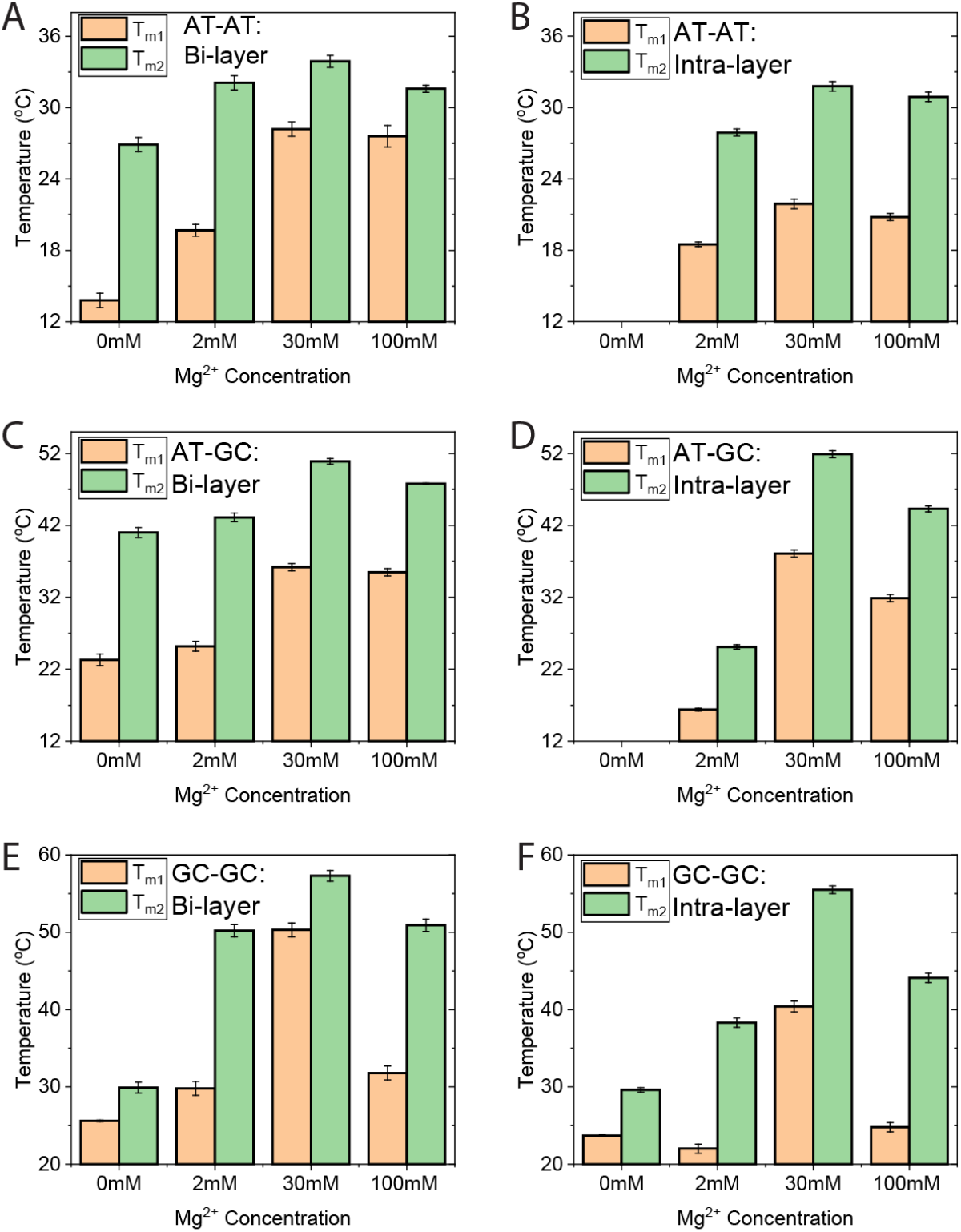
Thermal melting temperatures obtained from the double Hill function analysis in Fig. 3 and listed in Table 1. Note the general increase of their values with increasing Mg^2+^ concentration up to 30 mM, followed by a decrease at 100 mM. The y-axis scale of the charts has been started from an elevated temperature to better illustrate the contrast in the data.

## Discussion

Our results demonstrate that stacking of duplexes in a well-defined smectic layer structure, at a physiologically relevant GDNA concentration, provides a testbed (or “reference state”) to explore cation effects on ordering of duplexes within the layers and the thermal stability of that order. In the AT-AT and AT-GC solutions, the induction at lower temperatures of intra-layer positional ordering among the duplexes, as a consequence of introducing divalent cations, reduces the lateral separation between duplexes. To illustrate, the lateral separation is reduced from ∼36.1 Å in the smectic-A phase (determined by the center position of the diffuse wider-angle peak in Fig. 2A) to ∼33.5 Å in the smectic-B. This value is close to the result (∼30 Å) obtained in molecular dynamics (MD) simulations (6, 19) of DNA condensation. For each of the three constructs, the amplitude of the sharp wider-angle peak increases up to 30 mM added Mg^2+^, which corresponds to an increase in the amplitude of the density variation (Δ*ρ*_6_) that quantifies the degree of six-fold (hexagonal) positional ordering of the duplexes within the layers. The decrease in both Δ*ρρ*_6_ and thermal stability of the intralayer order at higher (100 mM) Mg^2+^ content, indicated by reduction in wider angle peak height in Fig. 2 and in melting temperatures (Table 1), is also consistent with MD simulations, where destabilization of the condensed state of DNA was attributed to a relatively uniform charge distribution developing at high divalent cation concentrations that would give rise to repulsive interactions (6), in contrast to the site-specific accumulation of ions in major grooves that was suggested to occur at lower ionic concentration.

By preparing samples with DNA concentration in the 260-265 mg/ml range, we targeted the transition between smectic-A and B phases observed for the AT-AT construct in the absence of divalent cations(29, 30). With this choice, the impact of Mg^2+^ on intralayer order should be easiest to detect, even at the low concentrations (∼ 1mM) that would be relevant *in vivo*. This strategy exposed condensation of DNA duplexes from fluid-to solid-like lateral packing (smectic-A to B transition) at Mg^2+^ concentration at least as low as 2 mM in solutions of the AT-AT and AT-GC GDNA constructs. These constructs also have weaker end-to-end base stacking interaction than the GC-GC construct (30), which already exhibits smectic-B ordering at 260-265 mg/ml DNA with no added Mg^2+^. Our parallel measurements on GDNA constructs with different terminal base pairings clearly reveal an interplay between the strength of end-to-end stacking interactions, which promote bilayer formation and inter-layer order, and side-to-side electrostatic interactions mediated by divalent cations, which act to stabilize intra-layer order.

To our knowledge, DNA condensation has only been demonstrated when cations with +3 or +4 valency were present, with the sole exception being the case where a 20-bp long duplex DNA condensed in the presence of divalent cations (in the absence of higher valency cations) when A-tracts, which adapt a distinct major groove structure, were introduced into the sequence (6). In that study, no condensation was observed for other sequences, including a construct with a random sequence. This ‘sequence enhanced major groove binding’ of ions was proposed as the mechanism that drives condensation of the A-tract sequence in the presence of Mg^2+^. Distinctly, we observe condensation with DNA constructs that have a random sequence in the presence of ∼2 mM Mg^2+^ concentration, and in the special case of GC-GC construct even in the absence of Mg^2+^ (just 150 mM Na^+^).

We have presented results of SAXS measurements on the effect of divalent cations on smectic liquid crystalline ordering in GDNA solutions with similar DNA and monovalent ion concentration and for three different terminal base pairings. When positively charged Mg^2+^ cations were introduced at low mM concentration into solutions of either AT-AT or AT-GC constructs in the bilayer smectic-A phase, the onset of lateral positional ordering (smectic-B phase) of the duplexes was observed, indicating a reduction of the repulsion between negatively charged phosphate backbones. For the GC-GC solution, initially in the smectic-B phase, introduction of Mg^2+^ reduced the repulsion between the negative phosphate backbones and further stabilized the lateral positional order. Increasing the Mg^2+^ concentration to 30 mM further enhanced the thermal stability of the smectic-B order, resulting in higher melting temperatures. These stabilizing effects have been confirmed by recent simulations and attributed to divalent cation accumulation in specific regions of the double helix structure, such as major grooves, and the alignment of these regions with phosphate backbones of neighboring helices. However, such correlations in charge localization are supposed to be reduced at higher Mg^2+^ concentration, where the ions are distributed in a more uniform “atmosphere”, mitigating attractive electrostatic interaction between helices. Our experimental results support this scenario. More generally, our work sheds light on the DNA condensation process, particularly when base-stacking or entropic considerations facilitate the condensation process beyond what is feasible with purely electrostatic interactions. These insights will provide better guidelines for molecular modeling efforts as they illustrate effective condensation at ionic conditions not observed previously.

## Materials And Methods

### GDNA Synthesis

Details of the GDNA synthesis process, sample loading process into thin-walled borosilicate capillaries, and the details of the SAXS/WAXS measurements carried out are described in previous publications (29, 30, 32). Briefly, the GDNA constructs, denoted as 48-20T-48, consist of two symmetric 48-base pair duplexes bridged by a single-strand gap of 20 Thymine bases (Fig.1A). These constructs are created by annealing a long strand (O1, 116 nt) with two short strands (O2 and O3, each 48 nt) which are complementary to the regions except the 20T sequence in the middle of O1. Polyacrylamide gel electrophoresis (PAGE) purified oligomers (O1, O2, and O3) were purchased from ExonanoRNA (Columbus, OH) or Genescript (Piscataway, NJ). The sequences of O1, O2, and O3 strands used to synthesize the GDNA constructs (AT-AT, AT-GC, and GC-GC) are given in (Supplementary Table S1). MgCl_2_ was added during the process of loading annealed GDNA solutions to the capillaries, yielding a final Mg^2+^ concentration of 2, 30, or 100 mM.

### Thermal Melting Analysis

In order to assign characteristic temperatures to the melting of the bilayer stacking or hexagonal intra-layer order, we first integrated the area under the associated peaks in Fig. 2 – namely, the first (fundamental) and third order small angle peaks (the latter barely visible in Fig.2 due to linear scale of the vertical axis) specifically associated with the bilayer stacking, and the sharp wider-angle peak associated with the intra-layer ordering. As described in more detail previously (30), the square root of the area scales with the magnitude of the density modulation or order parameter that characterizes the degree of positional order in each case. The areas under the SAXS peaks are calculated by fitting the data in each peak to a Gaussian function of *q*, and then integrating the fit results over *q* as shown in Supplementary Fig. S1. The square root of the resulting total integrated intensity under the relevant peaks is normalized with respect to the relevant maximum value and then plotted as a function of temperature in Fig. 3. The results were fit to single and double Hill functions of the form, 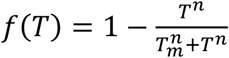 and 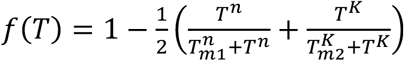, where *f*(*T*) is the average fraction of “bound” duplexes, *T*_*m*,_ *T*_*m1*,_ and *T*_*m2*_ are characteristic melting temperatures, and *n* and *k* are positive exponents. As noted in the caption to Fig. 3, in certain cases the double Hill function produces a kink in a region where the data is sparse, and which therefore may be artificial. With the tight constraints on synchrotron beam time, it was not feasible to perform measurements at higher temperature resolution.

## Acknowledgements

This study was supported by the National Science Foundation under grant DMR-1904167. The authors are particularly grateful to Ruipeng Li and Masa Fukuto for their assistance in performing the SAXS measurements on the CMS beamline (11-BM) at the National Synchrotron Light Source II, a U.S. Department of Energy (DOE) Office of Science User Facility operated for the DOE Office of Science by Brookhaven National Laboratory under Contract No. DE-SC0012704.

## Supplementary Data Available

Details of GDNA synthesis and SAXS measurements

## Notes

### Competing Interest Statement

The authors have declared no competing interest.

## References

1. Dove, A., Walton, R., Hamley, I., Mckeown, N., Robertson, N., Skinner, S., Cava, R., Cooper, A., Frs, C.G., Heeger, A., et al. (2012) Electrostatics of DNA compaction in viruses, bacteria and eukaryotes: functional insights and evolutionary perspective. Soft Matter, 8, 9285–9301.

2. Bloomfield, V.A. (1996) DNA condensation. Curr. Opin. Struct. Biol., 6, 334–341.

3. Bloomfield, V.A. (1997) DNA condensation by multivalent cations. Biopolymers, 44, 269–282.

4. Lipfert, J., Doniach, S., Das, R. and Herschlag, D. (2014) Understanding nucleic acid-ion interactions. Annu. Rev. Biochem., 83, 813–841.

5. Pelta, J., Livolant, F. and Sikorav, J.L. (1996) DNA aggregation induced by polyamines and cobalthexamine. J. Biol. Chem., 271, 5656–5662.

6. Srivastava, A., Timsina, R., Heo, S., Dewage, S.W., Kirmizialtin, S. and Qiu, X. (2020) Structure-guided DNA–DNA attraction mediated by divalent cations. Nucleic Acids Res., 48, 7018–7026.

7. Luan, B. and Aksimentiev, A. (2008) DNA Attraction in Monovalent and Divalent Electrolytes. J. Am. Chem. Soc., 130, 15754–15755.

8. Milo, R. and Phillips, R. (2015) Cell Biology by the Numbers First Edit. Milo, Ron; Phillips, R. (ed) Garland Science.

9. Tan, Z.J. and Chen, S.J. (2006) Nucleic acid helix stability: Effects of salt concentration, cation valence and size, and chain length. Biophys. J., 90, 1175–1190.

10. Anderson, C.F. and Record, M.T. (1995) Salt-nucleic acid interactions. Annu. Rev. Phys. Chem., 46, 657–700.

11. Korolev, N., Lyubartsev, A.P. and Nordenskiöld, L. (1998) Application of polyelectrolyte theories for analysis of DNA melting in the presence of Na+ and Mg2+ ions. Biophys. J., 75, 3041–3056.

12. Olmsted, M.C., Anderson, C.F. and Record, M.T. (1991) Importance of oligoelectrolyte end effects for the thermodynamics of conformational transitions of nucleic acid oligomers: A grand canonical Monte Carlo analysis. Biopolymers, 31, 1593–1604.

13. Hartwig, A. (2001) Role of magnesium in genomic stability. Mutat. Res., 475, 113–121.

14. Honig, B. and Nicholls, A. (1995) Classical Electrostatics in Biology and Chemistry. Science, 268, 1144–1149.

15. Kornyshev, A.A., Lee, D.J., Leikin, S. and Wynveen, A. (2007) Structure and interactions of biological helices. Rev. Mod. Phys., 79, 943–996.

16. Grochowski, P. and Trylska, J. (2008) Continuum molecular electrostatics, salt effects, and counterion binding—A review of the Poisson–Boltzmann theory and its modifications. Biopolymers, 89, 93–113.

17. Misra, V.K. and Draper, D.E. (1999) The Interpretation of Mg2+ Binding Isotherms for Nucleic Acids using Poisson-Boltzmann Theory. J. Mol. Biol., 294, 1135–1147.

18. He, W. and Kirmizialtin, S. (2020) Exploring Cation Mediated DNA Interactions Using Computer Simulations. Lect. Notes Bioeng., 10.1007/978-3-030-47705-9_6/FIGURES/4.

19. Yoo, J. and Aksimentiev, A. (2016) The structure and intermolecular forces of DNA condensates. Nucleic Acids Res., 44, 2036–2046.

20. Nakata, M., Zanchetta, G., Chapman, B.D., Jones, C.D., Cross, J.O., Pindak, R., Bellini, T. and Clark, N.A. (2007) End-to-end stacking and liquid crystal condensation of 6-to 20-base pair DNA duplexes. Science, 318, 1276–1279.

21. Bellini, T., Zanchetta, G., Fraccia, T.P., Cerbino, R., Tsai, E., Smith, G.P., Moran, M.J., Walba, D.M. and Clark, N.A. (2012) Liquid crystal self-assembly of random-sequence DNA oligomers. Proc. Natl. Acad. Sci. U. S. A., 109, 1110–1115.

22. Smith, G.P., Fraccia, T.P., Todisco, M., Zanchetta, G., Zhu, C., Hayden, E., Bellini, T. and Clark, N.A. (2018) Backbone-free duplex-stacked monomer nucleic acids exhibiting Watson–Crick selectivity. Proc. Natl. Acad. Sci. U. S. A., 115, E7658–E7664.

23. Davidson, M.W., Strzelecka, T.E. and Rill, R.L. (1988) Multiple Liquid Crystal Phases at High DNA Concentrations. Nature, 331, 457–460.

24. Fraccia, T.P., Smith, G.P., Bethge, L., Zanchetta, G., Nava, G., Klussmann, S., Clark, N.A. and Bellini, T. (2016) Liquid Crystal Ordering and Isotropic Gelation in Solutions of Four-Base-Long DNA Oligomers. ACS Nano, 10, 8508–8516.

25. Durand, D., Doucet, J. and Livolant, F. (1992) A study of the structure of highly concentrated phases of DNA by X-ray diffraction. J. Phys. II, 2, 1769–1783.

26. Onsager, L. (1949) The Effects of Shape on the Interaction of Colloidal Particles. Ann. N. Y. Acad. Sci., 51, 627–659.

27. Bolhuis, P. and Frenkel, D. (1997) Tracing the phase boundaries of hard spherocylinders. J. Chem. Phys., 106, 666–687.

28. Salamonczyk, M., Zhang, J., Portale, G., Zhu, C., Kentzinger, E., Gleeson, J.T.T., Jakli, A., De Michele, C., Dhont, J.K.G.K.G., Sprunt, S., et al. (2016) Smectic phase in suspensions of gapped DNA duplexes. Nat. Commun., 7, 13358.

29. Gyawali, P., Saha, R., Smith, G.P., Salamonczyk, M., Kharel, P., Basu, S., Li, R., Fukuto, M., Gleeson, J.T., Clark, N.A., et al. (2021) Mono- and bilayer smectic liquid crystal ordering in dense solutions of “gapped” DNA duplexes. Proc. Natl. Acad. Sci. U. S. A., 118, e2019996118.

30. Kodikara, S.G., Gyawali, P., Gleeson, J.T., Jakli, A., Sprunt, S. and Balci, H. (2023) Stability of End-to-End Base Stacking Interactions in Highly Concentrated DNA Solutions. Langmuir, 10.1021/acs.langmuir.3c00318.

31. Orellana, A.G. and De Michele, C. (2019) Free energy of conformational isomers: The case of gapped DNA duplexes. Eur. Phys. J. E 2019 426, 42.

32. Gyawali, P., Saha, R., Kodikara, S.G., Li, R., Fukuto, M., Gleeson, J.T., Smith, G.P., Clark, N.A., Jakli, A., Balci, H., et al. (2022) Smectic-B phase and temperature-driven smectic-B to -A transition in concentrated solutions of ‘gapped’ DNA. Phys. Rev. Res., 4, 033046.

33. Protozanova, E., Yakovchuk, P. and Frank-Kamenetskii, M.D. (2004) Stacked-unstacked equilibrium at the nick site of DNA. J. Mol. Biol., 342, 775–785.

34. Yakovchuk, P., Protozanova, E. and Frank-Kamenetskii, M.D. (2006) Base-stacking and base-pairing contributions into thermal stability of the DNA double helix. Nucleic Acids Res., 34, 564–574.

35. Petersheim, M. and Turner, D.H. (1983) Base-Stacking and Base-Pairing Contributions to Helix Stability: Thermodynamics of Double-Helix Formation with CCGG, CCGGp, CCGGAp, ACCGGp, CCGGUp, and ACCGGUp. Biochemistry, 22, 256–263.

36. Kilchherr, F., Wachauf, C., Pelz, B., Rief, M., Zacharias, M. and Dietz, H. (2016) Single-molecule dissection of stacking forces in DNA. Science, 353, aaf5508.

37. Mergny, J.L. and Lacroix, L. (2003) Analysis of Thermal Melting Curves. Oligonucleotides, 13, 515–537.

38. Parisiades, P. (2021) A review of the melting curves of transition metals at high pressures using static compression techniques. Crystals, 11, 416.

