## Supplementary Information for "Liquid Crystalline Layering and Divalent Cations Cooperatively Enhance DNA Condensation"

Sineth G. Kodikara[a], Prabesh Gyawali[a], James T. Gleeson [a], Antal Jakli [a,b], Samuel Sprunt[a,b,*], and Hamza Balci[a,*]
[a] Department of Physics, Kent State University, Kent, OH 44242, USA;
[b] Advanced Materials and Liquid Crystals Institute, Kent State University, Kent, OH 44242.

KEYWORDS: DNA Condensation, Gapped DNA, Smectic liquid crystal, Small angle x-ray scattering, Electrostatic Screening

**Thermal Melting Analysis**

**Figure S1:** To calculate the thermal melting temperatures, the areas under each peak are calculated. First, each peak is fitted to a Gaussian function as shown in the diagram. Two representative Gaussian fits are shown for the AT-AT, 30mM $Mg^{2+}$ construct at 7 °C for the peaks at $q_1$ (at left) and $q_w$ (at right). Open symbols are data points and solid lines are the Gaussian fits. The area under each peak is shaded. The base of the Gaussian represents the background.

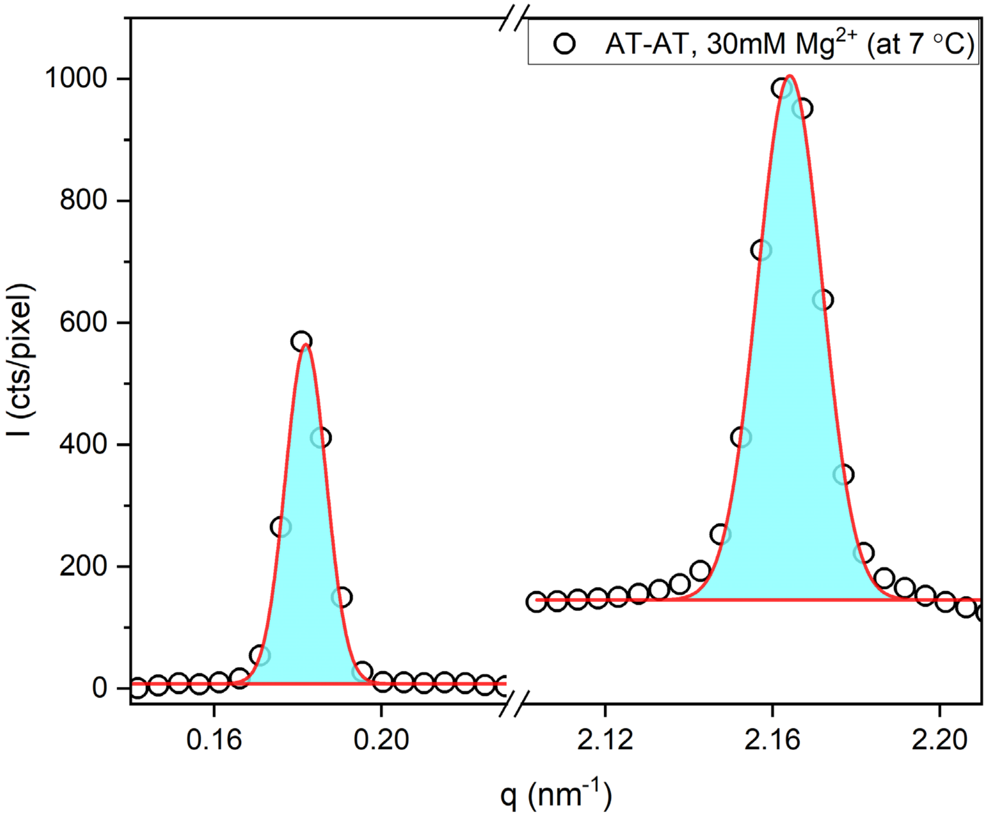

**DNA Sequences**

The O1 is the long strand (116 nt) in each case and O2 and O3 are the corresponding short strands (48 nt each).

**Table S1:** Sequences of oligonucleotides used for constructing three various GDNA samples.

| Construct | Strand | Sequence (5′ to 3′) | Length (nt) |
|---|---|---|---|
| AT-AT | O1 | ACAGATGCACATATCGAGGTGGACATCACTTACGCTGAGTACT TCGAATTTTTTTTTTTTTTTTTTTTTAAGCTTCATGAGTCGCATTC ACTACAGGTGGAGCTATACACGTAGACA | 116 |
| | O2 | TTCGAAGTACTCAGCGTAAGTGATGTCCACCTCGATATGTGCA TCTGT | 48 |
| | O3 | TGTCTACGTGTATAGCTCCACCTGTAGTGAATGCGACTCATGA AGCTT | 48 |
| AT-GC | O1 | ACAGATGCACATATCGAGGTGGACATCACTTACGCTGAGTACT TCGAATTTTTTTTTTTTTTTTTTTTTAAGCTTCATGAGTCGCATTC ACTACAGGTGGAGCTATACACGTAGACG | 116 |
| | O2 | TTCGAAGTACTCAGCGTAAGTGATGTCCACCTCGATATGTGCA TCTGT | 48 |
| | O3 | CGTCTACGTGTATAGCTCCACCTGTAGTGAATGCGACTCATGA AGCTT | 48 |
| GC-GC | O1 | GCAGATGCACATATCCAGGTGGACATCACTTACGCTGAGTACT TCGAATTTTTTTTTTTTTTTTTTTTTAAGCTTCATGAGTCGCATTC ACTACAGGTGGAGCTATACACGTAGACG | 116 |
| | O2 | TTCGAAGTACTCAGCGTAAGTGATGTCCACCTCGATATGTGCA TCTGC | 48 |
| | O3 | CGTCTACGTGTATAGCTCCACCTGTAGTGAATGCGACTCATGA AGCTT | 48 |

**DNA Sample Preparation**

The oligomers which were purified using polyacrylamide gel electrophoresis (PAGE) O1, O2, and O3 listed in the Supplementary Table S1 were obtained from ExonanoRNA or GenScript. To anneal the three strands, they were mixed in equal concentrations (10 µM) in a buffer containing 150 mM NaCl, 10 mM Tris-HCl (pH 7.5), and 0.1 mM ethylenediaminetetraacetic acid (EDTA). The mixture was heated to 90 °C for 10 minutes and then slowly cooled down to room temperature over several hours in a 1-gallon water heat bath.

The cooled-down annealed sample was filtered through a 50 kDa membrane (Amicon Ultra from Millipore) using centrifugation at 5,000 rpm for 15 minutes. The resulting supernatant, which had a volume of approximately 100 µL and a salt concentration of 150 mM NaCl, was collected. Deionized water was added to the supernatant, increasing the total volume by approximately five times and reducing the NaCl concentration to around 30 mM. This solution was then concentrated by passing it through a 10 kDa filter (Amicon Ultra from Millipore) and centrifuging it at 12,000 rpm for 12 minutes, maintaining the NaCl concentration at approximately 30 mM.

The concentration of GDNA in the concentrated supernatant was measured using a Nanodrop One instrument (Thermo Scientific) and was typically in the range of 80-100 mg/mL. Minor adjustments were made to the salt and GDNA concentrations of the supernatant to achieve a target DNA concentration of 260-265 mg/mL when the water evaporated from the open end of the capillary.

Before loading the GDNA samples, the capillaries were calibrated by adding known volumes of deionized water and measuring their height. The calibrated capillaries were then dried, and the supernatant GDNA samples were carefully pipetted into them. The initial concentration of $MgCl_2$ was calculated based on both the targeted $Mg^{2+}$ concentration and the final DNA concentration, and the calculated amount of $MgCl_2$ (from a 1 M stock) was added to each capillary.

The loaded capillaries were placed in a custom-made aluminum holder partially submerged in a water bath maintained at a temperature of 40-45 °C. As the water evaporated, the GDNA, NaCl, and $MgCl_2$ concentrations increased, while the solution height decreased. The capillary was removed from the bath and sealed with an inert epoxy when the desired height, and thus the desired GDNA and $MgCl_2$ concentrations, were achieved. The sealed capillaries were stored at 4 °C until the X-ray measurements were conducted, and the sample height was periodically checked to ensure the integrity of the seal.

**SAXS Measurements**

The beamline 11-BM at the National Synchrotron Light Source II (NSLS-II), was used to conduct the SAXS measurements. The incident X-ray energy delivered by a three-pole wiggler source was 17 keV, and the incident beam size at the sample was 0.2×0.2 $mm^2$. The SAXS detector was placed 3.0 m away from the sample. A commercially modified hot/cold stage (Instec HCS60) with Kapton film windows was used to mount the samples. The sample's temperature was maintained between 5 and 65°C (as the denaturation temperature of the GDNA duplexes of 48 bp is 77-80 °C depending on $Mg^{2+}$ concentration). The exposure time of the sample to the X-rays was 30 seconds at each step of the temperature. The background scattering was measured by recording the signal from a capillary filled with a buffer solution alone. This background signal was subtracted from the data obtained from the GDNA samples. To calibrate the scattering wave number (q) in the detector plane, a capillary filled with silver behenate powder was utilized. We did not observe any evidence of X-ray damage to the samples.
